# Model-free inference of evolution from allele frequency timeseries using permutation tests

**DOI:** 10.64898/2026.07.01.735864

**Authors:** Jason Bertram, Abigail Kushnir

## Abstract

Allele frequency (AF) timeseries allow us to directly observe the dynamics of evolution at a genetic level. However, extracting useful inferences from AF timeseries has proved difficult due to the model uncertainties and noisiness inherent in AF change at fine temporal scales. Here we present three new permutation tests — which do not assume a model of evolutionary change or a parametric statistical model — to detect AF timeseries features of evolutionary interest. The features identified by these approaches are: 1) any evolutionary change (as opposed to apparent change due to measurement error); 2) directional selection; 3) fluctuating selection with a propensity to change sign (negative autocorrelation). We are not aware of existing tests for features 1 and 3. Feature 2 is commonly tested using standard evolutionary models such as the Wright-Fisher; we show that the permutation approach has comparable statistical power. We apply our new approaches to AF timeseries data from D. melanogaster and D. pulex.

## 1 Introduction

Population genetic inference has historically focused on contemporaneous data such as fixed nucleotide differences [26] or the structure of segregating variation. It is becoming increasingly common for populations to be sequenced at multiple time points in evolve and resequence experiments [21, 1], natural populations [17, 14, 2, 20], as well as studies based on museum specimens [23] and ancient DNA [18]. Sequencing at multiple time points produces allele frequency (AF) timeseries — one timeseries for each measured polymorphic locus — which gives insight into evolutionary processes over much shorter timescales than contemporaneous data.

AF timeseries were the basis for some of the earliest inferences of natural selection [10]. Historically the essence of the problem has been to determine whether observed AF changes can be reasonably explained by a combination of random genetic drift and measurement error, or if it is necessary to invoke selection. A standard approach is to assume AFs are governed by a population genetic models like Wright-Fisher, Moran or the corresponding diffusion limit [4, 13, 18, 9, 15, 19, 24, 22, 7]. Henceforth we refer to these closely-related models collectively as “Wright-Fisher” for brevity. Since the effective population size *N* describing generational AF changes due to random genetic drift is often not known, these model-based approaches jointly infer *N* and the selection coefficient *s*, which is assumed to be constant over time for tractability.

Even for a single locus (a single AF timeseries), the inference procedures above can get quite mathematically involved and computationally demanding since we must effectively evaluate the probabilities of AF trajectories through time over many generations. A great deal of work has focused on these technical issues, even though it is not known whether simple models like Wright-Fisher accurately reflect biological reality in this context. A major justification for using Wright-Fisher is that many different models converge in their predictions in the diffusion limit [8], but this need not apply to the fine-grained dynamics captured in many AF timeseries.

Moreover, there is growing interest in characterizing the influence of fluctuating selection on genetic diversity [17, 25]. This is difficult to incorporate into the approaches described above since it is necessary to specify a model for the time-dependence of *s* and many models are *a priori* plausible. The addition of extra parameters would also amplify the computational challenge.

The problem is simpler if genetic drift can be neglected over the timescale of AF measurements. Then *s* can be estimated using regression [16, 17]. However, it is unlikely that drift can be ruled out *a priori* in most systems.

In other fields where timeseries are important, particularly econometrics and finance, the approach is typically to infer a generic model (often an auto-regressive stochastic process) from the timeseries rather than assume a mechanistic model governing the dynamics at the outset based on knowledge of the processes involved [12]. This approach is poorly suited to AF timeseries because it requires orders of magnitude more timepoints than is typically available, at least currently (∼ 10). Nevertheless, a more model-agnostic perspective could be useful in the analysis of AF timeseries.

Here we set out to develop AF timeseries inference methods that focus on identifying qualitative features of evolutionary interest rather than assuming a particular evolutionary model. This avoids the technical issues with evaluating trajectory probabilities under Wright-Fisher, while also giving insight into how much we gain from including the mechanistic details encoded in Wright-Fisher. Another aim of the present work is to relax the assumption of constant *s*, which is challenging to do in model-based approaches.

The basis of our approach is the permutation test. Permutation tests are non-parametric hypothesis tests based on resampling data in a manner that assumes a given null hypothesis is true [11]. A simple example is to test for a difference between two groups by permuting group identity in the data, where the null hypothesis is no group difference. In the case of AF timeseries, our permutation procedures will focus on the time dependence of measured AFs or AF changes. Being non-parametric has the obvious drawback of preventing estimation of common parameters like *s*. What we gain in return is: 1) the computational procedure is simple to describe and implement; 2) the tests are independent of evolutionary modeling assumptions such as Wright-Fisher or constant *s*.

Our approach is similar in spirit to studies based on calculating temporal autocovariance [6, 5] and among-locus timeseries divergence [3]. These approaches similarly do not rely on an evolutionary model, but leverage the fact that selection differs from neutral evolution in that it may have a preferred direction over time. A drawback of these approaches is reliance on combining AF information over many loci. For instance, in [5] the covariance between neighboring timepoints is computed across loci. As a result, only processes affecting the whole genome (or at least large parts of it) are captured. The methods developed here allow inference at a single locus, thereby allowing for much finer genomic resolution.

We develop three new permutation tests for the analysis of AF timeseries. Our first test, frequency permutation, checks if the timeseries is distinguishable from white noise — what we expect to observe if measurement error is the dominant contributor to AF changes. This test reveals that many empirical timeseries exhibit no detectable signals of evolution.

Our second test, sign permutation, checks for a consistent directional bias in AF changes, as would occur under directional selection. Surprisingly we find that sign permutation has similar power to detect selection as the influential model-based approach of [9], despite not being informed by a model of evolutionary dynamics.

Our third test, increment permutation, is similar to the second in that that it analyzes the directional structure of an AF timeseries, although this variant is less sensitive to directional selection and more sensitive to changes in direction (negative autocovariance). Unfortunately we find that this test is also sensitive to measurement error (which generates false positives), limiting its applicability to current datasets.

## 2 Methods

We assume that our data consists of a timeseries of the form

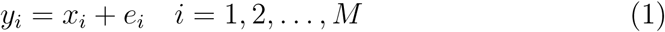

where *x_i_*is the AF in the population, *e_i_*is the measurement error and there are *M* timepoints in the timeseries. The index *i* labels measurements. Measurements are not necessarily taken in consecutive generations. Indeed the measurement times *t*_1_*, t*_2_*, …, t_M_* need not be evenly spaced, although for simplicity we will assume even spacing in our simulations. The increments corresponding to the observed AF timeseries are defined as Δ*y_i_*= *y_i_*_+1_ − *y_i_*for *i* = 1, 2*, …, M* − 1.

Since we will apply permutation tests to the *y_i_*, we do not require an explicit model for *x_i_* or *e_i_*. However, interpreting the test results does rely on assumed properties of these processes which will be described in the following sections.

### 2.1 Frequency permutation test

The frequency permutation (FP) test is based on permuting the order of AF observations. Intuitively, this tests if there is any non-random temporal structure in the time series, where a random (temporally unstructured) timeseries is represented by white noise. Expressed in terms of allele frequencies, the null hypothesis for this test is

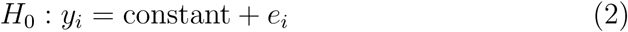

where the underlying AFs *x_i_* = constant are not evolving over time and measurement error *e_i_*is discrete white noise (the *e_i_*are independent and identically distributed (iid)).

*H*_0_ is false if (1) evolution causes the *x_i_*to change over time or (2) the *e_i_*differ from white noise. Regarding (1), any evolution, even pure genetic drift, is fundamentally different from white noise in that it causes cumulative AF changes. Consequently two AFs in the timeseries will tend to differ more the greater their time separation even if there is no trend in the timeseries (e.g. due to directional selection). We discuss (2) further below.

The FP test statistic is the mean increment magnitude

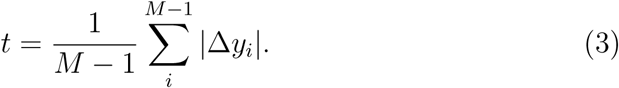

The FP test proceeds as follows. We repeatedly permute the order of the *y_i_*, and calculate *t* for each permutation. The distribution obtained in this way is used as the null distribution for *t*. This null distribution is then used to calculate a *p*-value for the observed *t*, denoted *t*_0_. Specifically, we calculate the one-sided *p*-value that the observed *t*_0_ is unusually small

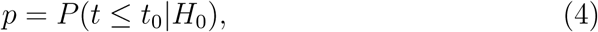

where the probability is calculated as the proportion of permuted trajectories satisfying *t* ≤ *t*_0_.

If *H*_0_ is false because the *x_i_* are evolving, then since permutation makes distant time points neighbors, the |Δ*y_i_*| are expected to increase (see (1) above). Hence *t*_0_ will tend to be unusually small compared to the null distribution for *t*.

Regarding the other way that *H*_0_ can fail (2 above), measurement error resembles white noise in that it is not uncorrelated over time and is not cumulative (unlike evolutionary change). However, Var[*e_i_*] may differ over time due to differences in sequencing and sampling effort. In most cases we do not expect this to generate false positives under Eq. (4) since the main effect is to conceal signals of evolution even more at timepoints with lower sampling effort. Hence, we interpret a rejection of *H*_0_ as indicating the presence of evolutionary AF change. We explore this idea further below with simulations.

### 2.2 Sign permutation test

The sign permutation (SP) test is based on permuting the signs of the increments Δ*y_i_*. Intuitively, this tests if there is non-random directionality in AF change. The null hypothesis is that positive and negative AF changes have equal probability

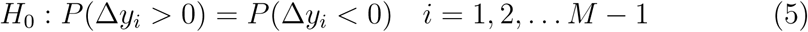

The alternative hypothesis if *H*_0_ is rejected is that either positive or negative changes are more likely. In our interpretation of the SP test we assume that the alternative hypothesis implies the presence of selection with a directional bias. In particular, we assume genetic drift does not produce a meaningful directional bias. We do not attempt to disentangle selection from migration, which could conceivably produce similar biases. Thus, expressed in evolutionary terms, SP tests for the presence of directional selection, where selection is interpreted in the broad sense of the combination of direct and indirect (linked) effects.

A particularly simple way to implement the SP test would be to use the number of positive increments (Δ*y_i_ >* 0) as the test statistic. Under the null hypothesis, this test statistic is binomially distributed with *M* − 1 trials and success probability 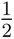. One drawback of this approach is that it gives equal weight to any magnitude of AF change (a large positive change is counted the same as a miniscule one). Consequently, we instead use the magnitude of the total displacement

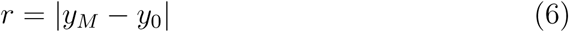

as the test statistic.

If there is a consistent directional component to the timeseries (e.g. a constant selection coefficient), the values of *r* generated by permutation will tend to be smaller than the observed *r*. The opposite occurs if the direction of the Δ*y_j_* tends to change.

The latter possibility is intriguing since it could in principle be used to directly test for the presence of fluctuating selection with negative temporal autocorrelation (e.g. as might occur with seasonal cycles). Unfortunately, negatively autocorrelated selection is hard to disentangle from measurement error which also induces negative correlations in neighboring increments (*e_i_*_+1_ − *e_i_*and *e_i_*_+2_ − *e_i_*_+1_ are negatively correlated because *e_i_*_1_ is shared).

For this reason we focus on detecting consistent directional change, which means using the *p*-value

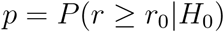

where *r*_0_ denotes the value of *r* in the observed (unpermuted) timeseries.

### 2.3 Increment permutation test

The increment permutation (IP) test is based on permuting the order of the increments Δ*y_i_*. Similar to FP, this tests if there is any non-random temporal structure in the series of AF changes. Our null hypothesis is

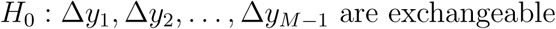

Exchangeability means that the joint probability distribution *P* (Δ*y*_1_, Δ*y*_2_*, …,* Δ*y_M−_*_1_) is invariant under permutations in the order of the sequence Δ*y*_1_, Δ*y*_2_*, …,* Δ*y_M−_*_1_. Intuitively, this means that the timeseries formed by permuting the Δ*y_i_* will have the same statistical properties as the observed timeseries.

In a simple random walk, the increments are iid, which implies exchangeability. Although random genetic drift is not a simple random walk since the magnitude of AF changes is frequency dependent, this frequency dependence has no appreciable effect on the IP test (Fig. S5).

The null hypothesis fails in two main ways: 1) the underlying AF changes Δ*x_i_* are not exchangeable; 2) Δ*y_i_*has a strong measurement error component — measurement error induces an order structure since it creates negative autocorrelations (see 2.2).

The test statistic we propose is the total deviation of a trajectory, defined as

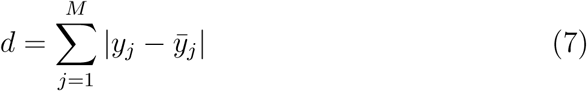

where *y̅* is the line connecting the start and end of the trajectory. Intuitively, *d* quantifies how far the AF timeseries strays from the shortest path connected its start and end.

The null distribution for *d* is computed by repeatedly permuting increments and calculating *d*. Similar to the SP test, we could test for the observed value of *d*, denoted *d*_0_, being either unusually small or large depending on the signal of interest. Unlike the SP test though, IP is ill-equipped to detect consistent directional selection since this form of selection is compatible with *H*_0_. On the other hand, IP is good at detecting negative autocorrelations, albeit with similar issues as SP regarding false negatives created by measurement error. Consequently, we use the *p*-value

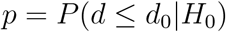

which corresponds to a test for *d*_0_ being unusually small.

### 2.4 Simulations

To analyze the tests above we simulate trajectories with a Wright-Fisher model. That is, we draw the frequency *x_t_*_+1_ in generation *t* + 1 from a binomial distribution with *N* trials and success probability *x_t_* + *s_t_x_t_*(1 − *x_t_*) where *s_t_* is the selection coefficient in generation *t*. We assume *s_t_* is normally distributed with mean *s* and variance *σ*^2^. Following Feder et al. [9], the simulated timeseries are either long (*T* = 1000) or short (*T* = 100) each measured at either *M* = 10 or *M* = 50 evenly spaced timepoints. Measurement error is incorporated by sampling *y_i_* from a binomial distribution with sample size *n* and success probability 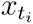 where *t_i_* is the time at which measurement *i* is taken. We consider no error, modest error (*n* = 1000) and high error (*n* = 100) scenarios.

### 2.5 Applications to empirical data

We apply the permutation tests developed here to two empirical datasets: 1) an evolve and resequence laboratory experiment with *D. melanogaster* [1] (*M* = 7 temporal measurements, 10 generations between measurements, ≈ 5 × 10^6^ loci) and 2) samples from a wild population of *D. pulex* [17] (*M* = 10 temporal measurements, approximately 4 generations between measurements, ≈ 9 × 10^5^ loci). Each locus represents a distinct AF timeseries.

Both datasets are used largely as is, with some minor preprocessing. The *D. melanogaster* data contains AF timeseries that go to fixation. We retain these trajectories but terminate them at the first instance of fixation, effectively resulting in a lower *M* for these timeseries. This dataset contains multiple replicate populations; for simplicity only the first replicate is analyzed. The *D. pulex* data contains some missing values: when these appear at the start/end of a timeseries, they are omitted (again reducing *M* for that timeseries) otherwise they are imputed as the mean of the neighboring AFs. For the *D. melanogaster* timeseries with *M* ≤ 7 we compute all possible permutations, resulting in exact *p*-values. Exhaustive permutation is computationally expensive for the *D. pulex* data with *M* = 10 (for the FP test there are a total of *M* ! permutations), and so we instead generate a sample of 10^4^ permutations. We include the unpermuted trajectory along with the set of permuted timeseries when computing *p*-values to avoid the pathological possibility of finding *p*-values exactly equal to zero.

As will be shown shortly, the combination of measurement error and modest *M* limits the ability to detect selection from a single locus in the empirical AF timeseries considered here. We therefore apply a windowing approach similar to ref [17] to combine information from multiple loci. For each window of 100 consecutive loci, for a given permutation test, we compute the mean *p*-value. We then compute the distribution of mean *p*-values obtained by randomly sampling 100 loci from across the genome. Since the latter distribution is normal, we use the corresponding best-fit normal cumulative density function to compute a *p*-value for each 100-consecutive-locus window. These *p*-values are then evaluated at a Bonferroni-corrected significance level to account for multiple testing. This approach leverages the fact that the signatures of evolution are shared by nearby loci (e.g. linked selection). It has the advantage of being straightforward and facilitating comparison with the results of [17].

## 3 Results

### 3.1 Simulations

The FP test reliably rejects the null hypothesis (white noise; no evolution) at the *α* = 0.05 significance level in the presence of genetic drift, directional selection, or fluctuating selection, provided that the relevant process is sufficiently strong compared to measurement error (Fig. 1). The statistical power (proportion of positives when *H*_0_ is false) also depends on the overall length of the timeseries and the number of measurements *M*. The data considered here covers a relatively short timespan (∼ 100) and a contain a modest number (∼ 10) of measurements. In this situation we need fairly strong drift (at most *N* ∼ 10^4^), a constant selection coefficient of *s >* 10*^−^*^3^ or a selection coefficient variance of *σ*^2^ *>* 10*^−^*^2^ to have power of at least 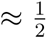. A high rate of false negatives is perhaps not surprising — AF timeseries with substantial measurement error, a modest number of timepoints, and a short overall evolutionary timespan are hard to distinguish from white noise. For completeness we also computed the receiver operating characteristic (ROC) for the FP test (Fig. S1), showing its performance as a function of the significance level *α*. The ROC also includes a scenario where measurement error varies strongly over time (assuming a Poisson distribution for *n*); it is shown that this causes reduction in the number of true positives rather creating false positives.

**Figure 1:**
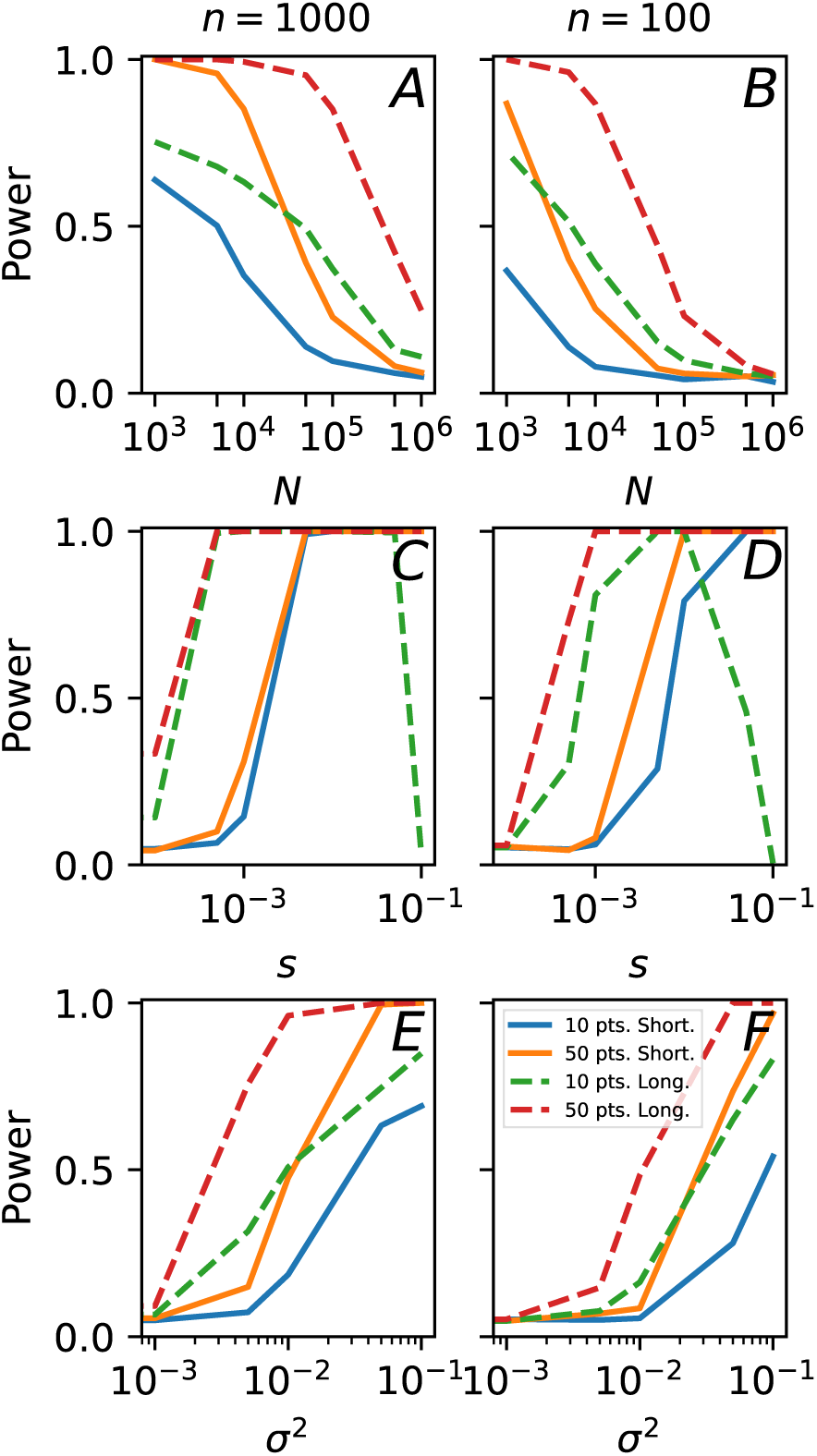
Power at the *α* = 0.05 significance level for the FP test as a function of population size *N*, selection coefficient *s* and selection coefficient variance *σ*^2^. Parameters when not specified by horizontal axis: *x*_0_ = 0.5, *σ*^2^ = 0, *s* = 0, *N* = 10^8^.

Turning to the SP test, the power to correctly identify directional selection (Fig. 2) depends on *Ns* which measures the strength of selection relative to drift. For low values *Ns* ∼ 1 the timeseries is dominated by drift and unsurprisingly the SP test fails to identify a signature of selection. Power greatly improves around *Ns >* 20. For very strong selective pressure, power actually starts to fall in long timeseries since fixation occurs. Measurement error greatly reduces power particularly for short timeseries (100 generations). The ROC for the SP test shows that measurement error only increases *p*-values (Fig. S2) — error does not generate false positives. These results are strikingly similar to the model-based approach of Feder, Kryazhimskiy, and Plotkin [9] (compare Figs. 2 and 5 in that reference).

**Figure 2:**
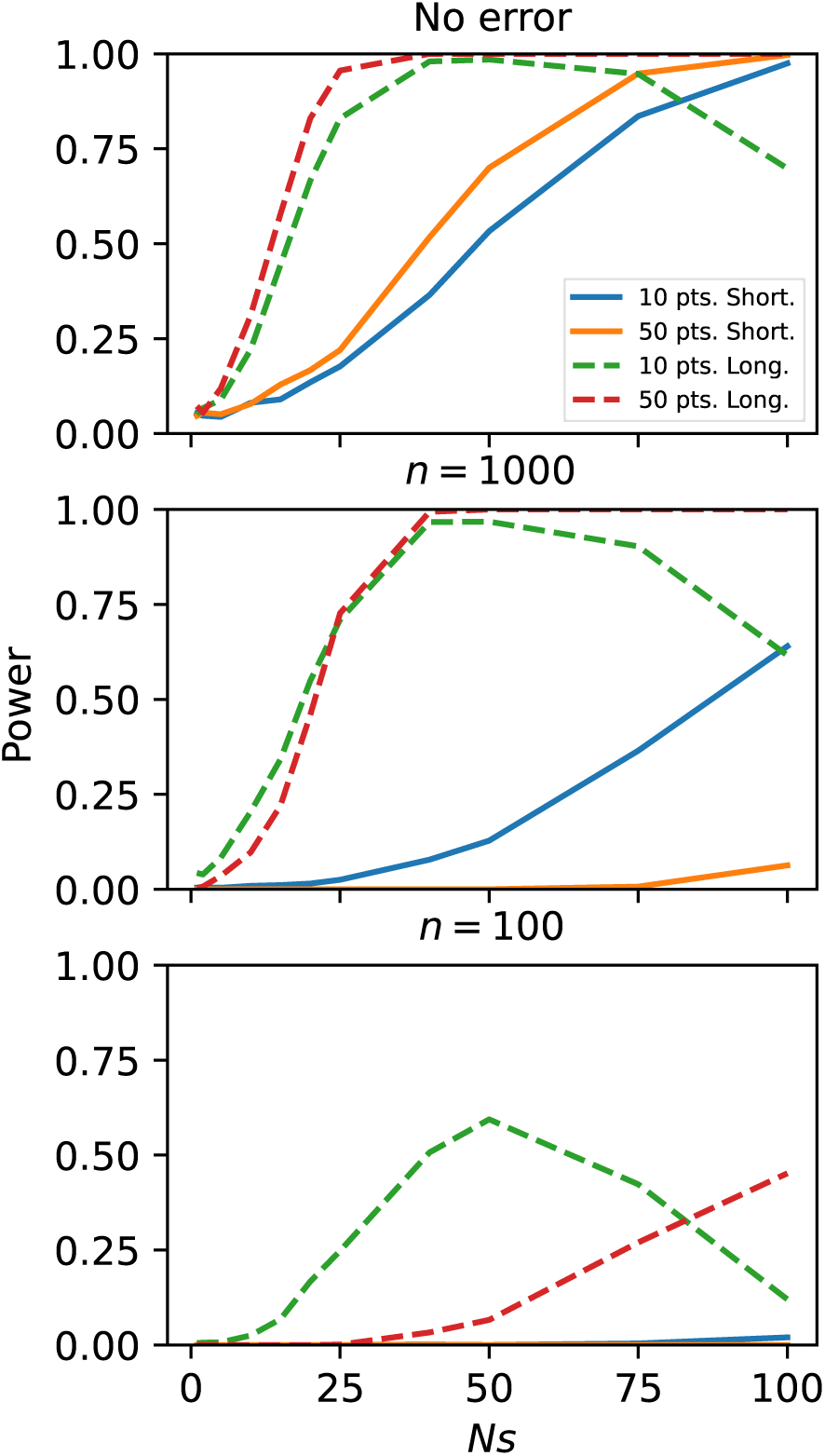
Power for the SP test at the *α* = 0.05 significance level vs the scaled selection strength *Ns*. Other parameters: *x*_0_ = 0.5, *σ*^2^ = 0, *N* = 10^4^.

Finally, we evaluate the ability of the IP test to detect negatively autocorrelated fluctuating selection. To do this, we simulate constant *s* within each interval separating measurements, but alternate the sign of *s* moving from one measurement interval to the next. In the absence of measurement error, the IP test correctly detects selection provided the strength of selection relative to drift is around *N* |*s*| *>* 100 (Fig. 3A). Introducing measurement error does not substantially reduce the rate of positives (Fig. 3B). However, as shown in Fig. 3C where there is no selection, measurement error greatly elevates the false positive rate of the IP test. Fig. 3C also shows that to be useful the IP test would need a sample size greater than *n* = 10^4^ in our binomial error model. Many datasets have much larger measurement error, including the datasets we consider below; hence, to avoid being mislead by false positives, we do not utilize the IP test in our empirical analysis.

**Figure 3:**
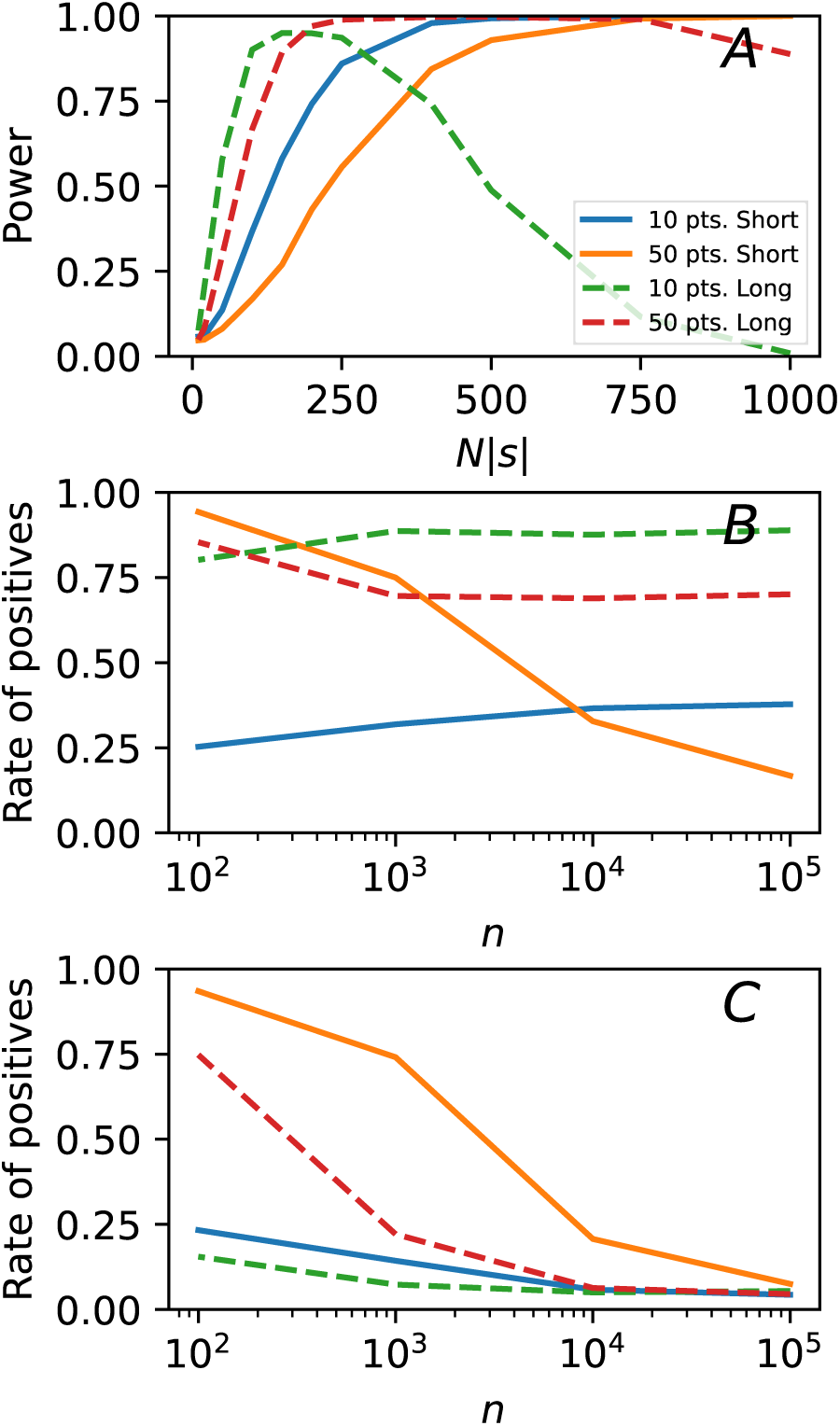
(A) Power for the IP test at the *α* = 0.05 significance level vs the scaled selection strength *N* |*s*|, where *s* is constant within each interval separating measurements, but alternates signs moving between measurement intervals. (B) Total rate of positives at the *α* = 0.05 significance level as a function of the binomial error sample size *n* for |*s*| = 0.01. (C) Same as B but with *s* = 0, so all positives are false. Other parameters: *x*_0_ = 0.5, *σ*^2^ = 0, *N* = 10^4^.

### 3.2 Application to empirical AF timeseries

We now apply our permutation tests to two whole-genome AF timeseries datasets. Our simulation results show that the permutation tests work in principle, although their power is fairly weak in short timeseries with *M* = 10 measurements, which is the simulation scenario representative of the datasets considered here and most datasets currently available. The windowing procedure described in 2.5 combines the permutation test results across groups of 100 consecutive loci and allows us to identify genomic regions undergoing any detectable evolution (FP test) or direction selection (SP test).

Fig. 4 shows the *D. pulex* results [17]. Even after windowing, there is strikingly little signal of evolution according to the FP test. Only a few scattered regions are genome-wide significant for directional selection according to the SP test. Of course, failure to reject the null hypothesis does not imply that it is true. There could be more directional selection (SP test) or evolutionary change at large (FP test) present in this dataset with our tests simply lacking power to detect it. It is nevertheless striking to contrast these *D. pulex* results with the *D. melanogaster* results (Fig. 5). For the latter, the entire genome shows a comfortable rejection of the white noise null hypothesis (FP test) and strong signals of directional selection (SP test).

**Figure 4:**
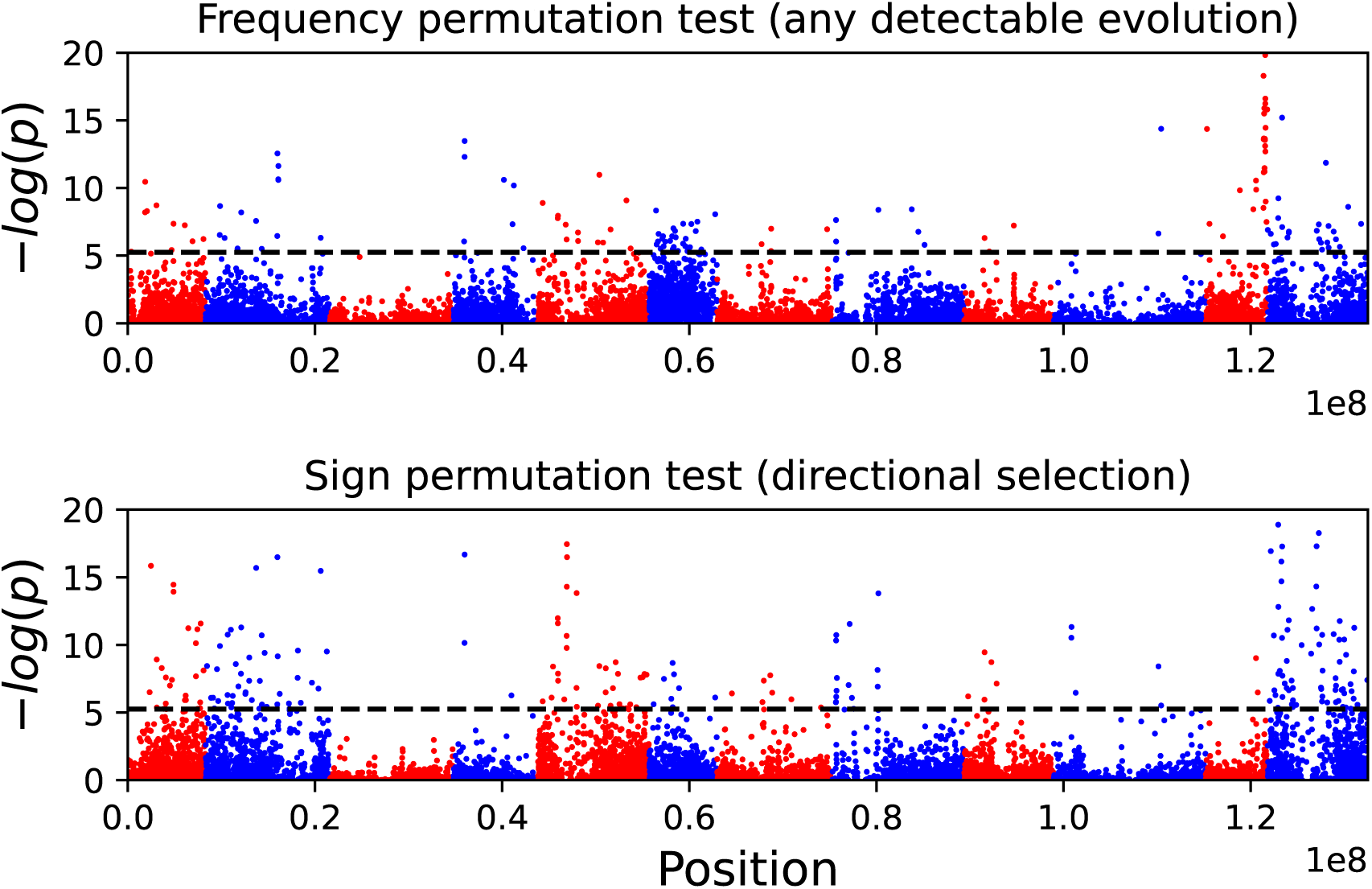
Windowed *p*-values (100 consecutive loci) for genome-wide AF timeseries data in D. pulex. Horizontal line shows Bonferonni-corrected *α*. Large parts of the genome show little detectable evolution under the FP test, and likewise for signatures of directional selection that are genome-wide significant under the SP test. Colors alternate between chromosomes.

**Figure 5:**
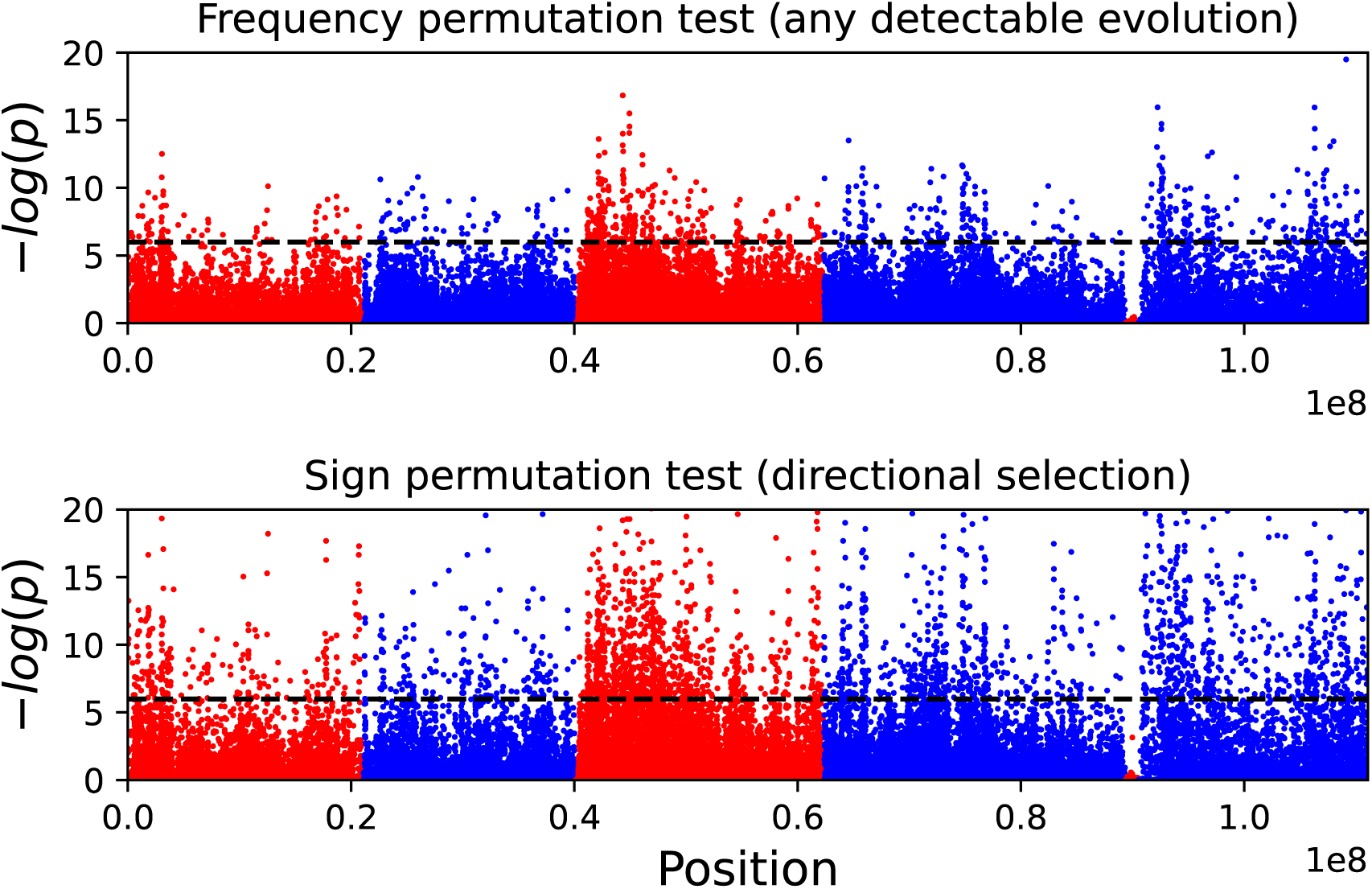
Windowed *p*-values (100 consecutive loci) for genome-wide AF timeseries data in D. melanogaster. Horizontal line shows Bonferonni-corrected *α*. Throughout the genome there are clear signatures of evolution according to the FP test, as well as strong signatures of directional selection. Colors alternate between chromosomes.

## 4 Discussion

We have presented three new permutation tests for inferring evolution from AF timeseries, respectively the FP test, the SP test and the IP test. The SP test addresses the common task of identifying selection in its simplest (i.e. directional) form, whereas the FP and IP tests are able to identify signals that are not currently identified by existing AF timeseries methods (respectively, any evolution and negatively autocorrelated selection). The applicability of the IP test is constrained by measurement error, which introduces false positives. The magnitude of measurement error is too great in the empirical data considered here for the IP test to be reliable.

The FP test can check how much evolutionary signal there is in an AF timeseries with relatively little effort. The FP test assumes white noise as its null hypothesis. This might seem strange since typically we take it for granted that observable AF evolution is occurring even over the modest duration (∼ 10 − 100 generations) of most temporally-resolved studies. However, in the presence of measurement noise, it cannot be taken for granted that a signal of evolution is observable in AF timeseries.

Indeed, we find weak signal of evolution in the *D. pulex* data of Lynch et al. [17] (Fig. 4), presumably due to the strength of measurement error rather than true AF stasis. Lynch et al. argue that random genetic drift is negligible over the timescale of their study (∼ 40 generations). Based on this, they attribute AF change entirely to selection, finding weak directional selection coefficients of order 10*^−^*^3^ and comparatively large temporal selection coefficient variance of order 10*^−^*^2^. These numbers are annual totals amounting to approximately 4 generations of evolution — hence the corresponding per-generation parameters *s* and *σ*^2^ in our analysis are likely even smaller (depending on the intra-annual structure of selection). The sample-size *n* is around 30 *< n <* 100.

Overall, our findings are consistent with Lynch et al. [17]. Namely, we expect evolution to be essentially undetectable by the single-locus FP test for effective population sizes above ∼ 10^4^ (Fig. 1B), for constant selection coefficients below ∼ 10*^−^*^3^ (Fig. 1D) or for selection coefficient variance below ∼ 10*^−^*^2^. Our SP test simulation results are harder to compare since the variance-effective population size *N* needed to compare *Ns* is not known. The windowed SP and FP analysis does nevertheless identify regions that are genome-wide significant, and these are broadly consistent with each other and with the genome-wide analysis in Lynch et al. [17] — for instance, chromosomes 3 and 4 show noticeably fewer genome-wide significant hits (Fig. 4). The *D. melanogaster* data in contrast shows a vast number of genome-wide significant hits across the genome (Fig. 5), consistent with the finding of large linkage blocks under strong selection by Barghi, Hermisson, and Schlötterer [1]. This signal is readily detected despite the fact that the *D. melanogaster* timeseries contain fewer measurements (*M* = 7 vs *M* = 10). The difference with the *D. pulex* study [17] might be partly due to larger sample sizes (typically *>* 100) or more generations of evolution between measurements (10 vs 4), although it seems unlikely that this is the whole story since these differences are modest. Genetic drift is likely much stronger in the laboratory *D. melanogaster* population as the census population size is only *N* ∼ 500. However, drift cannot not explain the SP test results. Given the preceding points, the SP test results indicate selection with a stronger directional component affecting a higher proportion of the genome compared with the *D. pulex* study [17].

Permutation tests are frequently used in evolutionary genetics, along with the closely-related bootstrapping and cross-validation resampling procedures [11]. To pick one salient example, Buffalo and Coop [5] apply their covariance approach to permuted data from Bergland et al. [2]. Thus, the novelty of our approach is not the mere fact of using permutation, but rather the particular choices of which variables are permuted and of test statistic, allowing AF features of evolutionary interest to be identified. We do not claim these choices are exhaustive — after all, perhaps the greatest feature of permutation tests is their flexibility.

We find it remarkable that the SP test performs on par with the likelihood approach of Feder, Kryazhimskiy, and Plotkin [9] (Fig. 2). That is, explicitly computing trajectory likelihoods under Wright-Fisher dynamics is approximately as informative about the presence of directional selection (in terms of statistical power) as checking for consistent directionality via sign permutation. Parametric models can go further than our permutation tests and estimate the intensity of selection, which is obviously important for evaluating evolutionary hypotheses. Nevertheless, we hope that due to their simplicity and relative freedom from dynamical assumptions, the permutation tests proposed here will be useful for exploratory work, and will help to inform the development and choice of parametric models.

## Data availability

Code used to generate simulation figures can be accessed at github.com/jasonbertram/trajectory_permuation.

## Acknowledgements

We thank Joanna Masel for valuable input on the role of measurement error.

## Supplement

**Figure S1:**
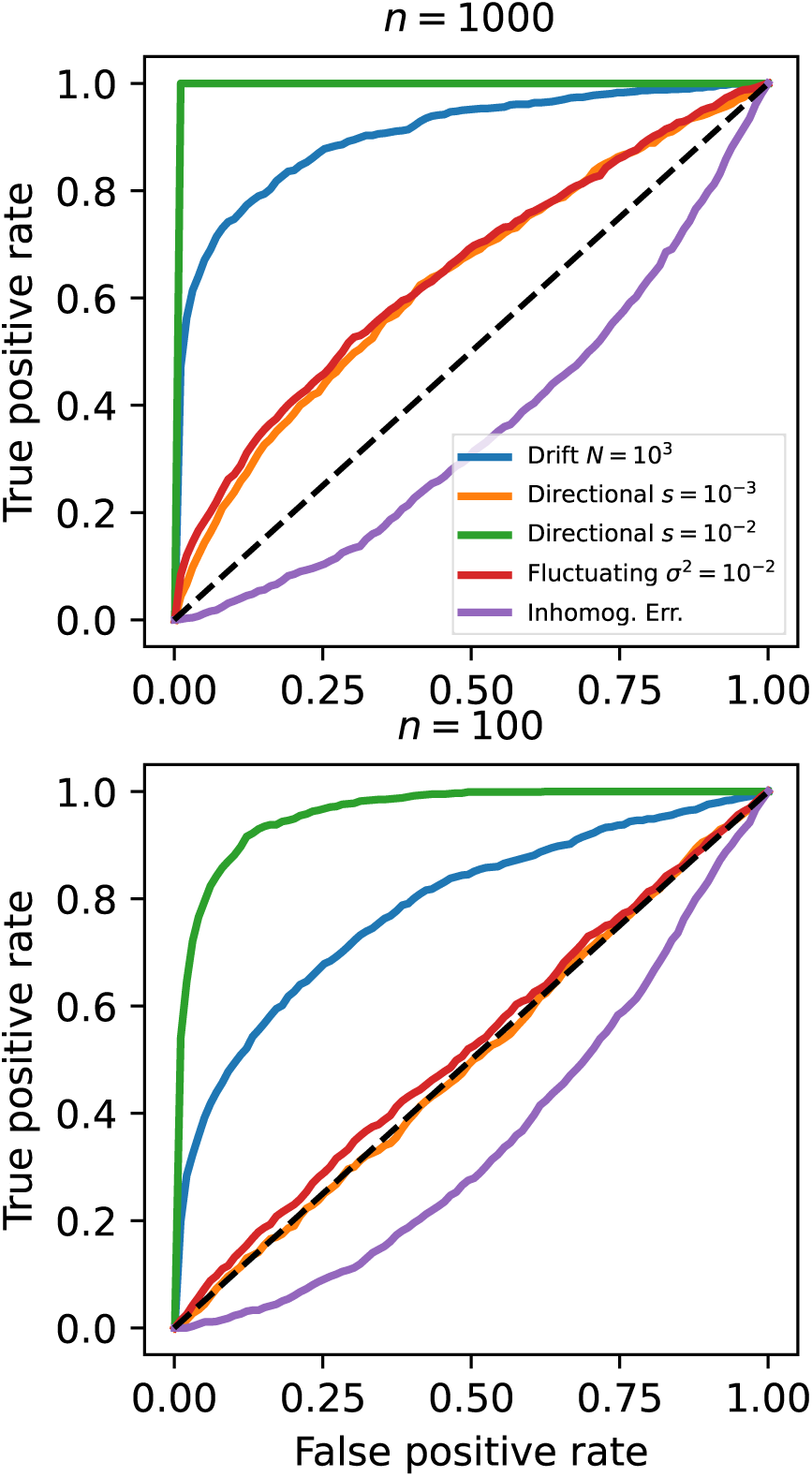
ROC for the FP test applied to Wright-Fisher simulations (10 measurements taken in 10 generation intervals). Six scenarios are considered: drift alone, directional selection with constant *s* (two different values), fluctuating selection with constant *σ*^2^ (two different values), and a high-variance inhomogeneous error scenario where the measurement error at each timestep is drawn from a Poisson distribution with mean *n*. Default parameters when unspecified: *N* = 10^10^, *s* = 0 and *σ*^2^ = 0.

**Figure S2:**
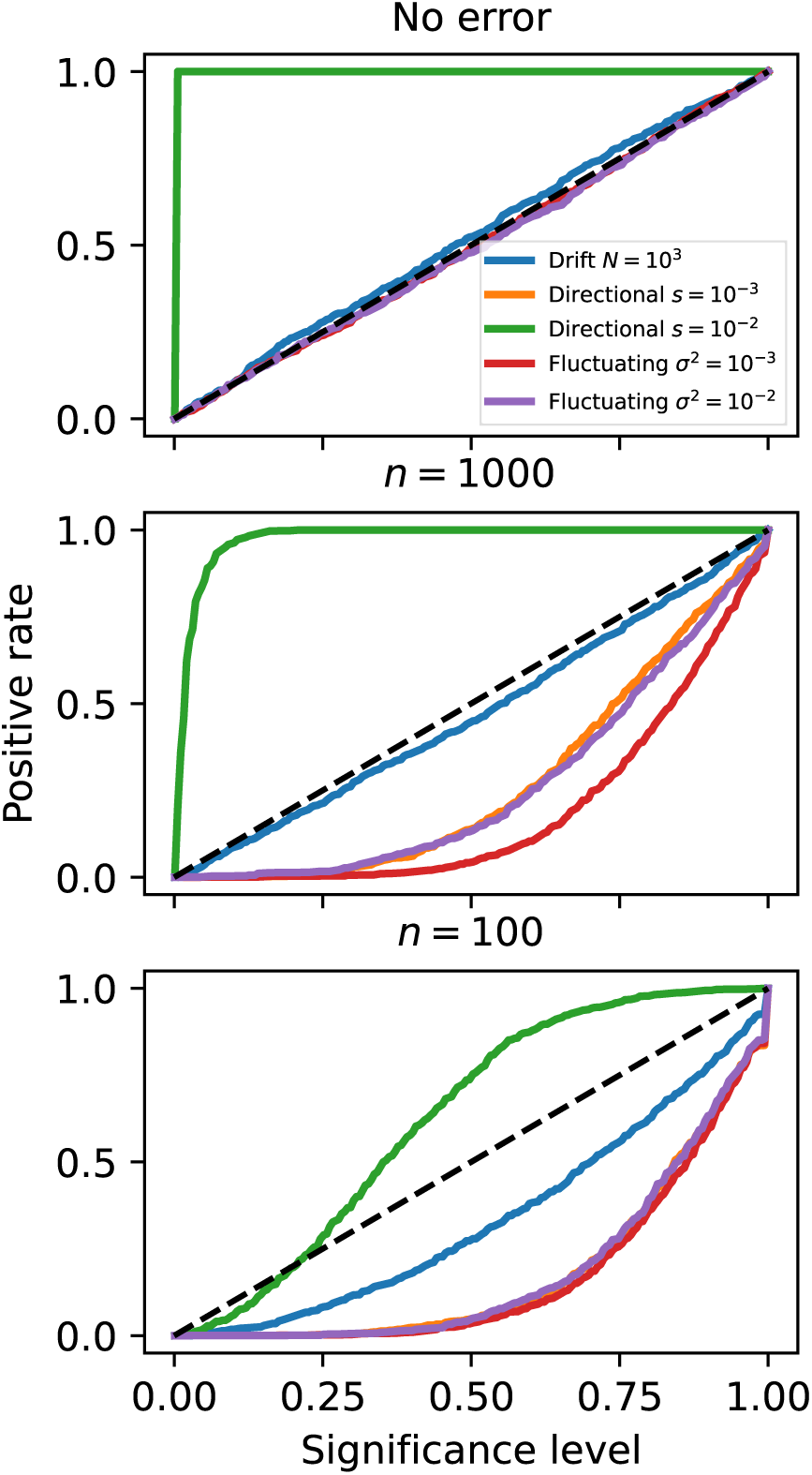
ROC for the SP test applied to Wright-Fisher simulations (10 measurements taken in 10 generation intervals). Measurement error increases *p*-values thereby reducing power.

**Figure S3:**
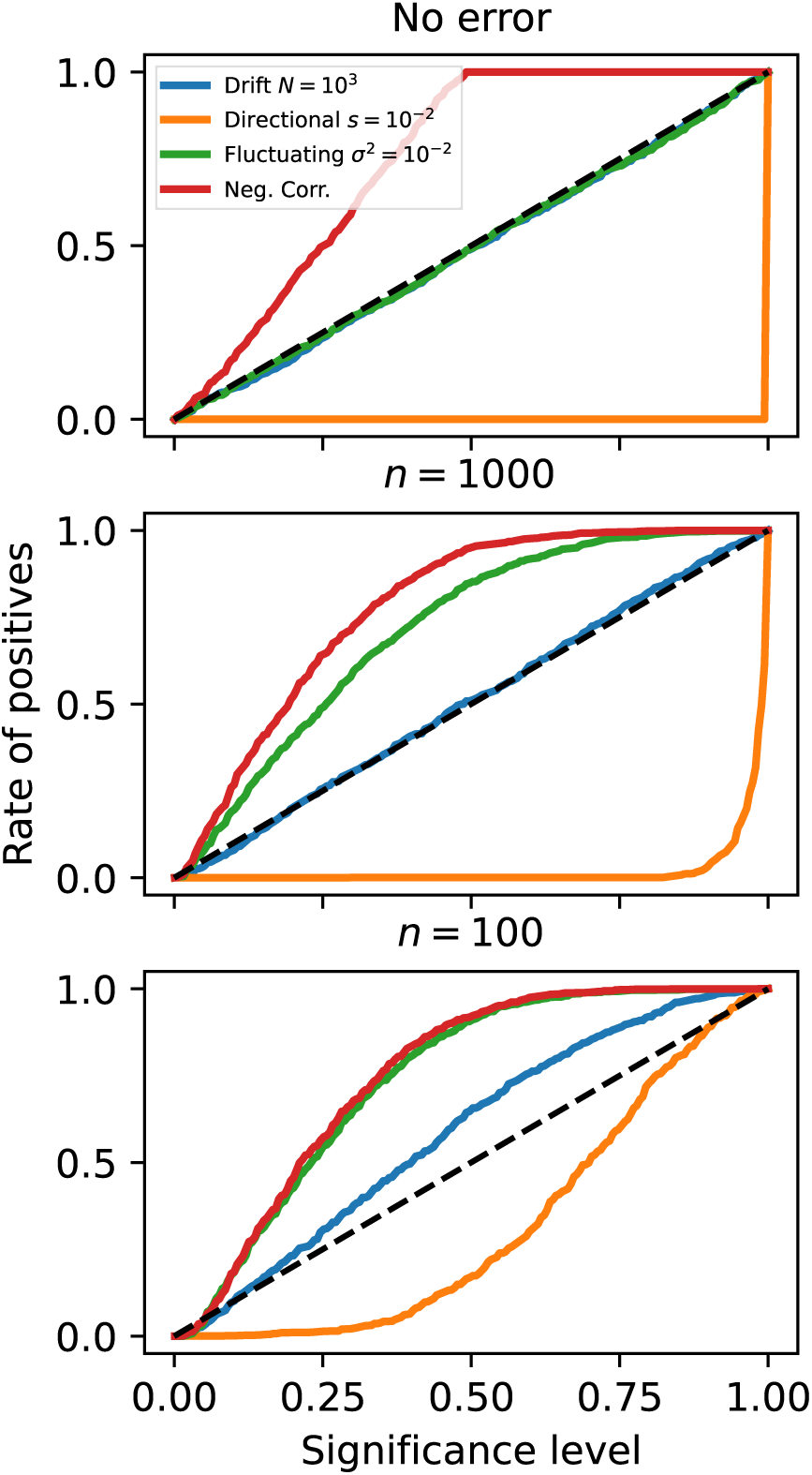
ROC for the SP test but using the alternate *p*-value of *p* = *P* (*r <*≤ *r*_0_|*H*_0_). Wright-Fisher simulations are the same as Fig. S2, but an alternating selection negative correlation scenario is now added (see IP results in 3.1).

**Figure S4:**
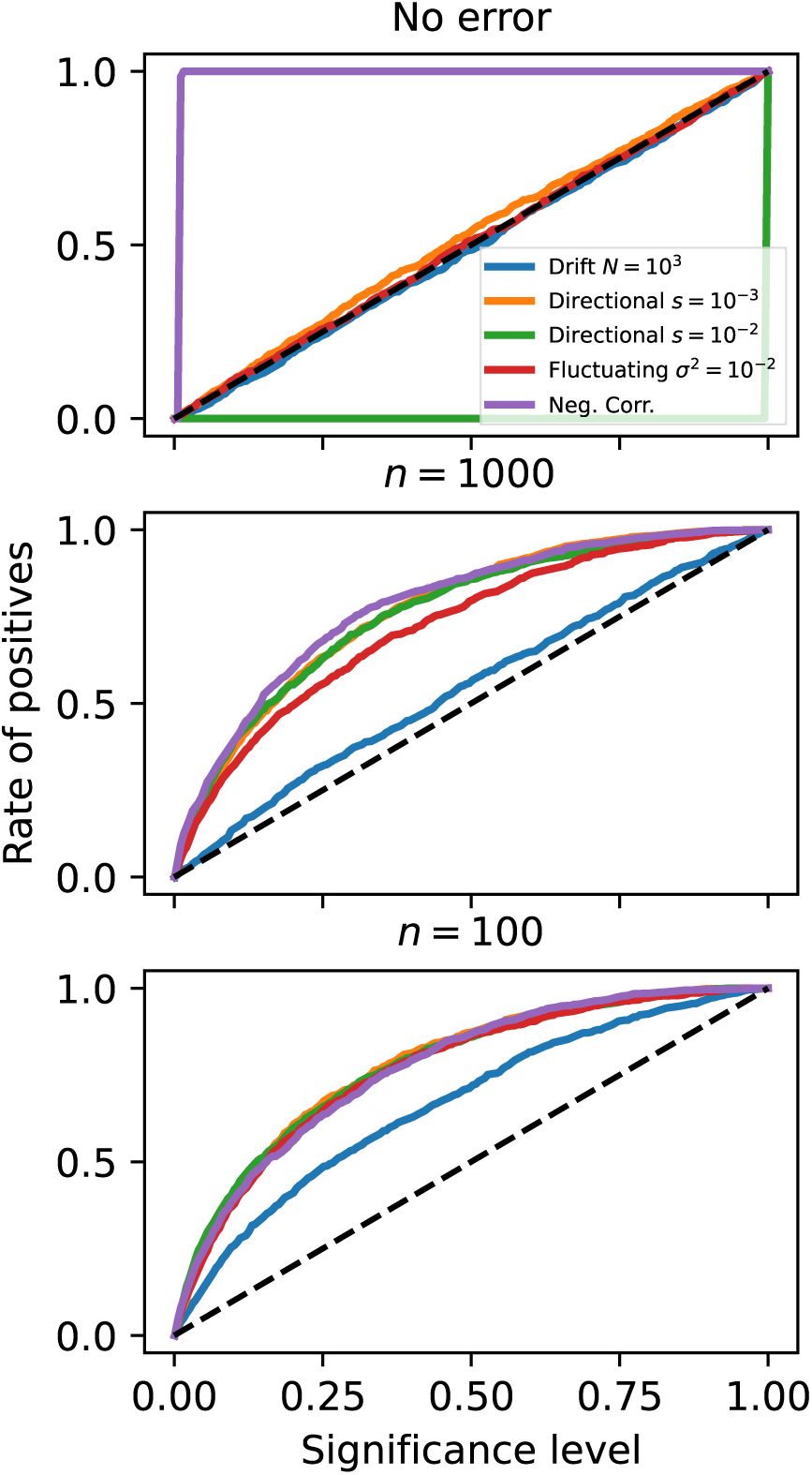
ROC for the IP test. Wright-Fisher simulations are the same as Fig. S2, but an alternating selection negative correlation scenario is now added (see IP results in 3.1). The power of the IP test to detect alternating selection is far superior to the SP test, although measurement error causes many false positives in all simulated scenarios.

**Figure S5:**
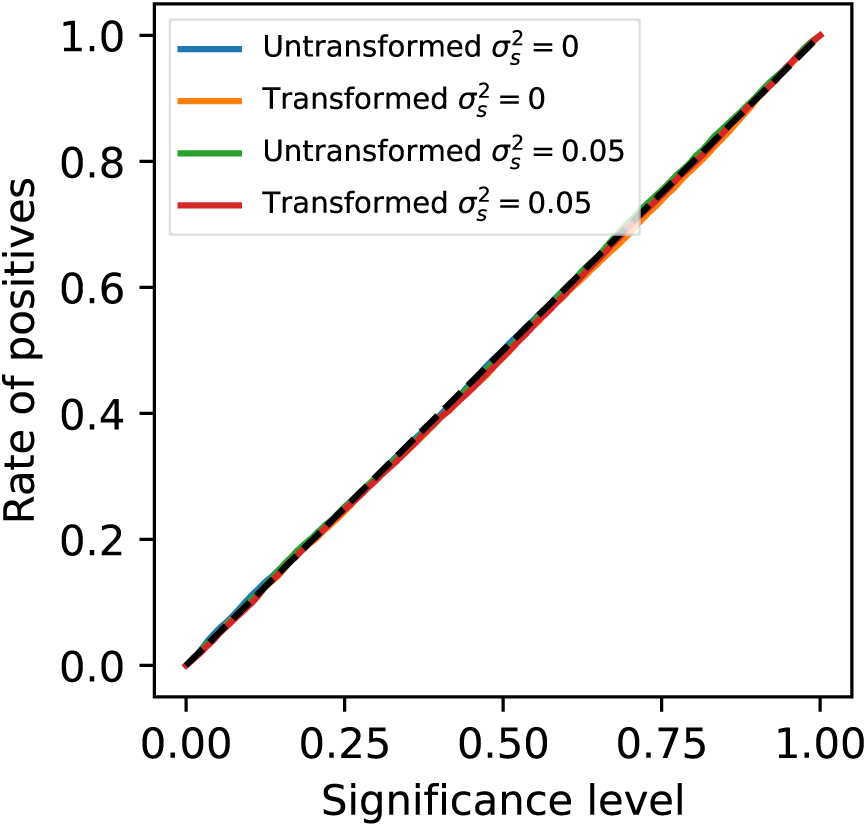
The increment permutation test rejects the null hypothesis at a rate indistinguishable from the significance level under a pure drift Wright-Fisher process 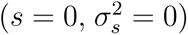 and a Wright-Fisher process with uncorrelated fluctuating selection 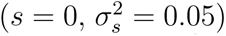, regardless of the use of the variance-stabilizing transformation 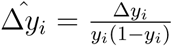. Other parameters: *N* = 10^3^, *x*_0_ = 0.2. 10 measurements taken in 10 generation intervals).

